# Prevalence of *Mycoplasmopsis agassizii* across wild and captive Mediterranean tortoises

**DOI:** 10.64898/2026.03.11.710774

**Authors:** Marta Canós-Burguete, Andrés Giménez, Albert Martínez-Silvestre, Joan Budó, Rachel E. Marschang, Beatriz Sánchez-Ferreiro, Roberto C. Rodríguez-Caro, Eva Graciá

## Abstract

*Mycoplasmopsis* [*Mycoplasma*] *agassizii* is one of the principal pathogens associated with upper respiratory tract disease (URTD) in tortoises, yet its epidemiology in European wild chelonian populations remains poorly understood. The pathogen has been linked to population declines in some wild tortoise populations and is frequently detected in captive tortoises, where infections may persist subclinically and prolonged contact can facilitate transmission. In this context, the pet trade and the release or escape of captive individuals represent potential pathways for pathogen exchange between captive and wild populations. We assessed the presence and prevalence of *M. agassizii* in wild Mediterranean tortoises in Spain and compared infection patterns with captive populations. A total of 259 tortoises were sampled between 2020 and 2025, including spur thighed tortoises (*Testudo graeca*; 127 wild; 63 captive) and Hermann’s tortoises (*Testudo hermanni*; 46 wild; 23 captive). Detection of *M. agassizii* was performed using PCR. The pathogen was detected in both species, but prevalence patterns differed markedly between captivity status and species. High prevalence was consistently observed in captive individuals of both species. In contrast, wild populations showed species-specific patterns: *T. graeca* exhibited very low or absent prevalence across wild populations, whereas *T. hermanni* showed comparatively higher prevalence in the wild. These results provide the first baseline assessment of *M. agassizii* occurrence in Mediterranean tortoises in Spain and highlight the importance of incorporating pathogen surveillance into conservation and management strategies for European chelonian populations.

## Introduction

Chelonians are among the most threatened vertebrate groups worldwide, with population declines primarily driven by habitat loss, global trade, overexploitation and climate change (Luiselli et al., 2016; Rodríguez-Caro et al. 2023; Chen et al. 2025; TCC 2025). Although infectious diseases are not considered major global drivers of decline for most turtle species, disease outbreaks can exacerbate demographic pressures in certain systems affecting chelonian population dynamics (Jacobson et al. 2014; Iglesias et al. 2015; Adamovicz et al. 2020; Graciá et al. 2020; TCC 2025). Their ectothermic physiology and long-life histories make chelonians particularly sensitive to environmental stressors such as thermal fluctuations and global trade, that may also influence disease susceptibility and amplify the impacts of pathogens through synergistic interactions with anthropogenic pressures (Altizer et al. 2003; Rios and Zimmerman 2015; Drake et al. 2019; Rush et al. 2021; Rodríguez-Caro et al. 2023, 2025; Di Nicola et al. 2025). Within this context, *Mycoplasmopsis* [*Mycoplasma*] *agassizii* has been recognized as one of the principal pathogens affecting chelonian health, with documented impacts in several species and rising concern for conservation (Brown et al. 1999; Jacobson et al. 2014; Ballouard et al. 2021; Louro et al. 2025).

*M. agassizii* is a primary etiological agent of upper respiratory tract disease (URTD) and is associated with morbidity and mortality in some wild and captive tortoise populations (Brown et al. 1999; Soares et al. 2004; Jacobson et al. 2014; Berry et al. 2015; Galosi et al. 2023). Clinical signs include rhinitis (from serous to mucopurulent nasal discharge), ocular discharge, conjunctivitis, and palpebral edema (Jacobson et al. 2014). Nevertheless, the response of tortoise species to *M. agassizii* infections may vary, with disparities in susceptibility, immune strategy and clinical manifestation (Berry et al., 2015; Weitzman et al. 2017; Sandmeier et al., 2023). Evidence from wild tortoise populations in the USA indicates that the severity of URTD may be influenced by environmental stressors such as drought or temperature extremes, which can elevate bacterial loads and increase morbidity and mortality risk (Sandmeier et al. 2009; Weitzman et al. 2017). In Europe, *M. agassizii* infections have been reported predominantly in captive settings, where close contact among individuals facilitates pathogen transmission (Origgi et al. 2004; Soares et al. 2004; Salinas et al. 2011b; Kolesnik et al. 2017; Galosi et al. 2023; Louro et al. 2025). Coinfections with other pathogens, such as herpesviruses or picornaviruses, have also been documented, adding complexity in the assessment of the health risks of infection to chelonians (Salinas et al. 2011b; Kolesnik et al. 2017; Adamovicz et al. 2020). Despite decades of research, the epidemiology of *M. agassizii* in European chelonians remains poorly understood, not only in terms of its occurrence in wild populations but also regarding the potential impacts it may exert on their population dynamics (Soares et al., 2004; Salinas et al., 2011b; Lecis et al., 2013; Jacobson et al., 2014; Kolesnik et al., 2017; Hidalgo-Vila et al., 2020; Ballouard et al. 2021; Marenzoni et al. 2022; Galosi et al. 2023; Louro et al. 2025).

The trade in Mediterranean tortoises is widespread, with large numbers of Hermann’s tortoises (*Testudo hermanni*) and spur-thighed tortoises (*Testudo graeca*) being sold in European and other international markets (Auliya et al. 2016; Graciá et al. 2020; Segura et al. 2020; Shearer and Türkozan 2024). Research on wild populations of *T. graeca* has shown that the presence of tortoises from captivity is relatively frequent, as shown by morphological assessments and genetic analyses that identify released or escaped captive individuals (Graciá et al. 2011; Salinas et al. 2011a; Semaha et al. 2024). Furthermore, the last native population of *T. hermanni* in the Iberian Peninsula has been reported to host non-native *Testudo* species (e.g. *T. marginata*), as well as other exotic chelonian species (e.g. *Centrochelys sulcata*), in addition to non-autochthonous intraspecific lineages of *T. hermanni*, highlighting the permeability of wild populations to human-mediated introductions (Pfau et al. 2020). Such intentional or accidental releases increase the likelihood of pathogen spillover from captive to wild populations (Aiello et al. 2014; Warne and Chaber 2023). In this context, *M. agassizii* infections have been reported in both captive and wild *Testudo* across Europe (Soares et al. 2004; Lecis et al. 2013; Ballouard et al. 2021; Galosi et al. 2023) and in the Iberian Peninsula (Salinas et al. 2011b; Martinez-Silvestre et al. 2022; Louro et al. 2025). This pathogen has also been detected in wild *T. graeca* in Morocco (Mathes 2003). Despite these reports, data on wild populations of Mediterranean tortoises in Spain remain scarce. On this basis, we hypothesise that infection prevalence would be higher in captive individuals than in wild populations, reflecting the role of captive environments in facilitating pathogen transmission and maintenance. Here, we assess the presence and prevalence of *M. agassizii* in wild populations of Mediterranean tortoises (*T. hermanni* and *T. graeca*) in Spain and compare these findings with infection levels in captive individuals of recovery centers to provide baseline epidemiological data that can inform future risk assessments and conservation actions.

## Methods

### Study species and sampling design

The spur-thighed tortoise (*Testudo graeca*) and the Hermann’s tortoise (*Testudo hermanni*) are the only native terrestrial chelonian species currently occurring in Spain, although their distributions are highly fragmented (Graciá et al. 2020). In the Iberian Peninsula, *T. graeca* occurs naturally in a geographically restricted area in the south-eastern region, specifically in the provinces of Murcia and Almería, and has also established a human-introduced population in the south-western Doñana National Park (Graciá et al. 2017a, b). In contrast, the native distribution of *T. hermanni* is currently limited to a single natural population in northeastern Spain, located in the Serra de l’Albera (Catalonia). Additional mainland populations of *T. hermanni* in Catalonia, including the Massís de la Collserola, Massís del Garraf, Massís dels Ports, Serra del Montsant, Serra de Llaberia, Pantà del Gaià and the Ebro Delta, originate from recient reintroduction programs, with further introduced populations reported in the Valencian Community (Pfau et al. 2020; Bertolero et al. 2020). In the Balearic Islands, both *T. graeca* and *T. hermanni* have been historically introduced to Mallorca, where they show an allopatric distribution, while *T. hermanni* has also been historically introduced to Menorca (Mateo et al. 2011, Pinya 2011; Bertolero and Pretus 2015, Graciá et al. 2017a). This complex biogeographical context, characterized by a mosaic of native and introduced populations, provides a relevant framework to assess pathogen presence and prevalence across captivity status and species, and to explore potential differences in host–pathogen dynamics between *T. graeca* and *T. hermanni*.

In this study, we evaluated 190 individual *T. graeca* and 69 *T. hermanni* to assess the presence of *M. agassizii* in Spain (Fig. 1). We sampled 127 wild and 63 captive *T. graeca* individuals. Wild tortoises were surveyed in Almería, Doñana and Mallorca, whereas captilve tortoises were surveyed at wildlife recovery centers (WRC) of Mallorca, Santa Faz (Alicante), and El Valle (Murcia). In addition, 46 *T. hermanni* were sampled from wild populations in Catalonia and Mallorca, and 23 captive *T. hermanni* from recovery centres for reptiles and tortoises (CRARC and CRT de l’Albera). During fieldwork in Mallorca, two *T. hermanni* individuals were encountered and sampled within areas defined a priori as *T. graeca* sampling sites, likely due to human release.

**Fig. 1.**
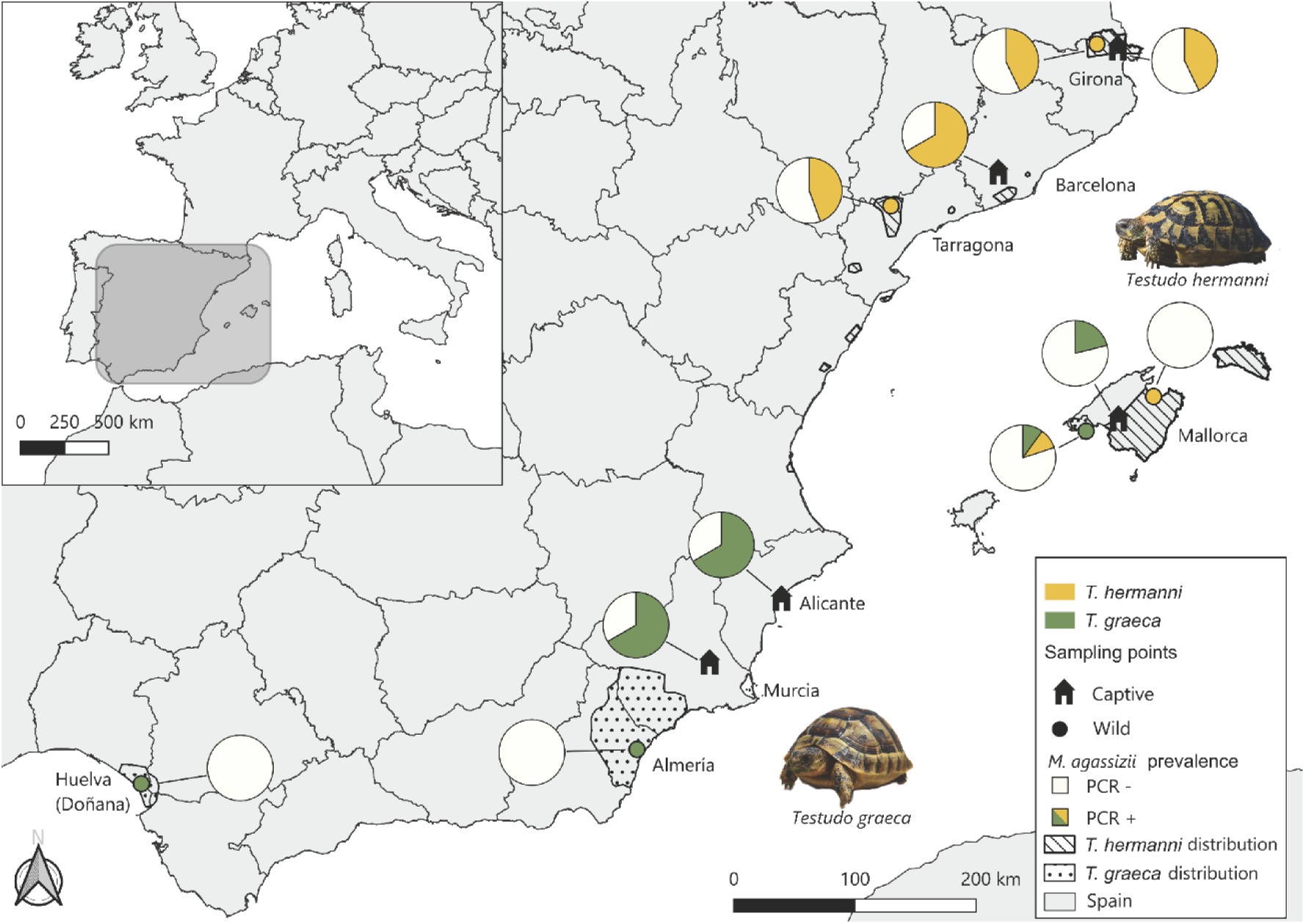
Geographic distribution of *Mycoplasmopsis agassizii* prevalence across Mediterranean tortoise populations in Spain. Pie charts indicate PCR results at each sampling location, filled segments denote PCR-positive individuals and open segments PCR-negative individuals, with green representing *T. graeca* and yellow *T. hermanni*. In Mallorca, one *T. hermanni* individual testing positive for *M. agassizii* was detected within a *T. graeca* population. Shaded areas indicate the distribution ranges of both species.

### Clinical assessment and molecular detection

During field surveys, tortoises encountered along transects were captured for individual marking and biological sampling. Wild and captive *T. hermanni* were randomly selected for inclusion and showed no clinical signs at the time of sampling. In *T. graeca*, individuals were also randomly selected from both wild and captive populations; most were asymptomatic, although 14% of captive individuals of WRC of Alicante and Murcia and one wild individual from Mallorca exhibited poor body condition and oral mucosal pallor. To minimize health risks and prevent cross-contamination, all individuals were handled using disposable medical gloves, which were changed or disinfected between animals (Martínez-Silvestre et al. 2023). Oronasal epithelial and mucosal samples were obtained by swabbing the throat, tongue, and palate. Swabs were preserved in ethanol, stored frozen, and subsequently analyzed at a specialized veterinary diagnostic laboratory (LABOKLIN GmbH & Co. KG, Bad Kissingen, Germany) using PCR assays for the detection of *M. agassizii* as described previously (Kolesnik et al., 2017). After arrival at the laboratory, samples were processed within 24 h. Briefly, swabs were soaked in 750 µl of lysis buffer (Roche Diagnostics, Mannheim, Germany) supplemented with 75 µl proteinase K (500 µg/ml; Carl Roth GmbH & Co. KG, Karlsruhe, Germany) and incubated for 20 min at 65°C. DNA and RNA were extracted from 200 µl of the lysis buffer using a commercial kit (MagNA Pure 96 DNA and viral NA small volume kit, Roche) according to the manufacturer’s instructions. Prepared DNA and RNA were stored at 4°C for 1 wk and then at −18°C. Samples were tested for the presence of *M. agassizii* and related *Mycoplasmopsis* spp. using the PCR targeting the V3 variable region of the 16S rRNA gene described by Brown et al. (1995) as described previously (Faulhaber et al., 2025). Each PCR plate included negative and positive controls as well as an extraction control (DNA Process Control Detection Kit, Roche Diagnostics GmbH, Mannheim, Germany) in each sample to assess nucleic acid extraction efficiency and PCR inhibition. Presence and size of the PCR product was assessed by capillary electrophoresis (QIAxcel Advanced system, Qiagen GmbH, Hilden, Germany). The expected product size was 576 bp.

### Statistical analyses

To assess factors associated with the presence and prevalence of *M. agassizii* in Mediterranean tortoises, we fitted generalized linear mixed models (GLMMs) with a binomial error distribution and logit link. Infection status (PCR positive/negative) was modelled as a function of captivity status (wild vs. captive), species, and their interaction, with sampling region included as a random intercept to account for spatial clustering. We compared all candidate models using AICc. Estimated marginal means and pairwise post hoc comparisons were then obtained, and marginal and conditional R² values were calculated to quantify the variance explained by fixed and random effects. All analyses were conducted in R Project.

## Results

*Mycoplasmopsis* [*Mycoplasma*] *agassizii* was detected in both *Testudo graeca* and *T. hermanni*, but prevalence varied markedly across species and between wild and captive status (Table 1, Figure 2). All *T. graeca* individuals presenting clinical signs at the time of sampling tested PCR-positive for *M. agassizii*. Captive *T. graeca* exhibited high infection levels (67% in WRC of Alicante and Murcia, 21% in WRC of Mallorca; Table 1), whereas wild populations showed almost no evidence of infection: animals from both Almería and Doñana tested negative, and only a single wild individual from Mallorca was positive (13%; Table 1). In *T. hermanni*, captive individuals also showed high prevalence (67% in CRARC, 43% in Albera CRT; Table 1), while wild populations displayed contrasting regional patterns, with high prevalence in Catalonia (43% Girona, 44% Tarragona; Table 1) but low infection rates in Mallorca (14%; Table 1). Nevertheless, the only PCR-positive *T. hermanni* on the island corresponded to a released individual in a *T. graeca* locality (Figure 1).

**Fig. 2.**
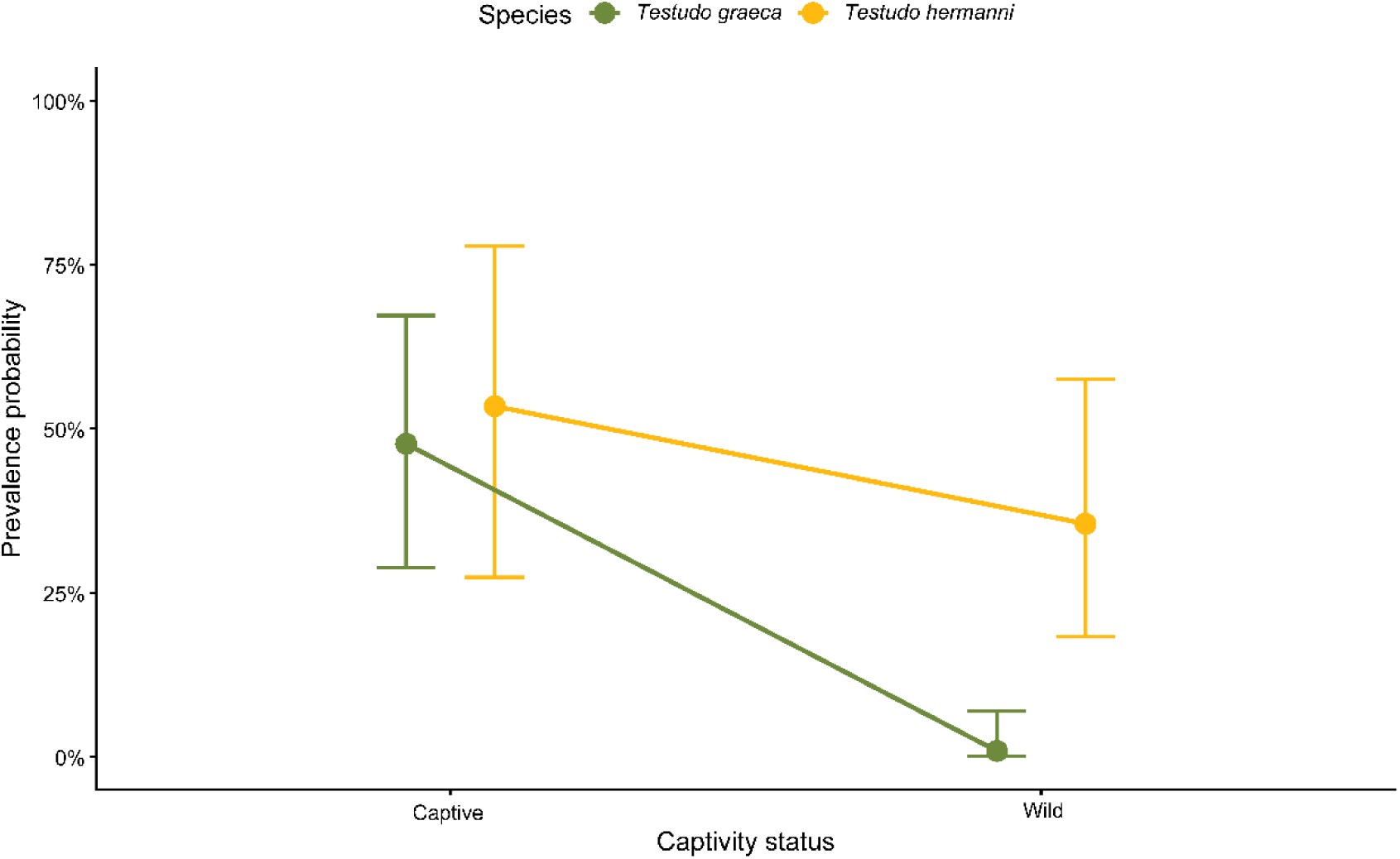
Predicted probability of *Mycoplasmopsis agassizii* infection in *Testudo graeca* and *Testudo hermanni* under captive and wild conditions, estimated from the best-supported generalized linear mixed model including a captivity status x species interaction and sampling region as a random effect. Points represent marginal means and error bars indicate 95% confidence intervals, back-transformed from the logit scale.

**Table 1.** Prevalence of *Mycoplasmopsis agassizii* across different populations of Mediterranean tortoises in Spain.

| **Captivity status** | **Specie** | **Sampling region** | **N** | **Positive (%)** |
| --- | --- | --- | --- | --- |
| Captive | *T. graeca* | WRC Alicante | 15 | 10 (67%) |
|  |  | WRC Murcia | 15 | 10 (67%) |
|  |  | WRC Mallorca | 33 | 7 (21%) |
|  |  | **Total** | **63** | **27 (43%)** |
| Captive | *T. hermanni* | CRT Girona | 14 | 6 (43%) |
|  |  | WRC Barcelona (CRARC) | 9 | 6 (67%) |
|  |  | **Total** | **23** | **12 (52%)** |
| Wild | *T. graeca* | Almería | 107 | 0 (0%) |
|  |  | Mallorca | 8 | 1 (13%) |
|  |  | Huelva (Doñana) | 12 | 0 (0%) |
|  |  | **Total** | **127** | **1 (0.007%)** |
| Wild | *T. hermanni* | Mallorca | 7 | 1 (14%) |
|  |  | Girona | 21 | 9 (43%) |
|  |  | Tarragona | 18 | 8 (44%) |
|  |  | **Total** | **46** | **18 (39%)** |

Model selection supported the effect of captivity status, species, and an interaction between these two variables (AICc weight = 0.84; Table 2). Estimated means derived from the GLMM indicated that in *T. graeca*, predicted prevalence dropped sharply from high levels in captivity (0.48; 95% CI: 0.29–0.67) to near zero in wild populations (0.009; 95% CI: 0.001–0.069). In contrast, *T. hermanni* showed consistently high prevalence, with similar infection probabilities in captivity (0.49; CI: 0.27–0.78) and the wild (0.36; CI: 0.18–0.58) (Figure 2). Post-hoc comparisons show that wild *T. graeca* had significantly lower prevalence than any other species-captivity status combination (*p* < 0.005, Holm-adjusted). The random effect of sampling region showed moderate variance (SD = 0.52), and the GLMM explained a substantial proportion of variation (R² marginal = 0.591; R² conditional = 0.622).

**Table 2.**
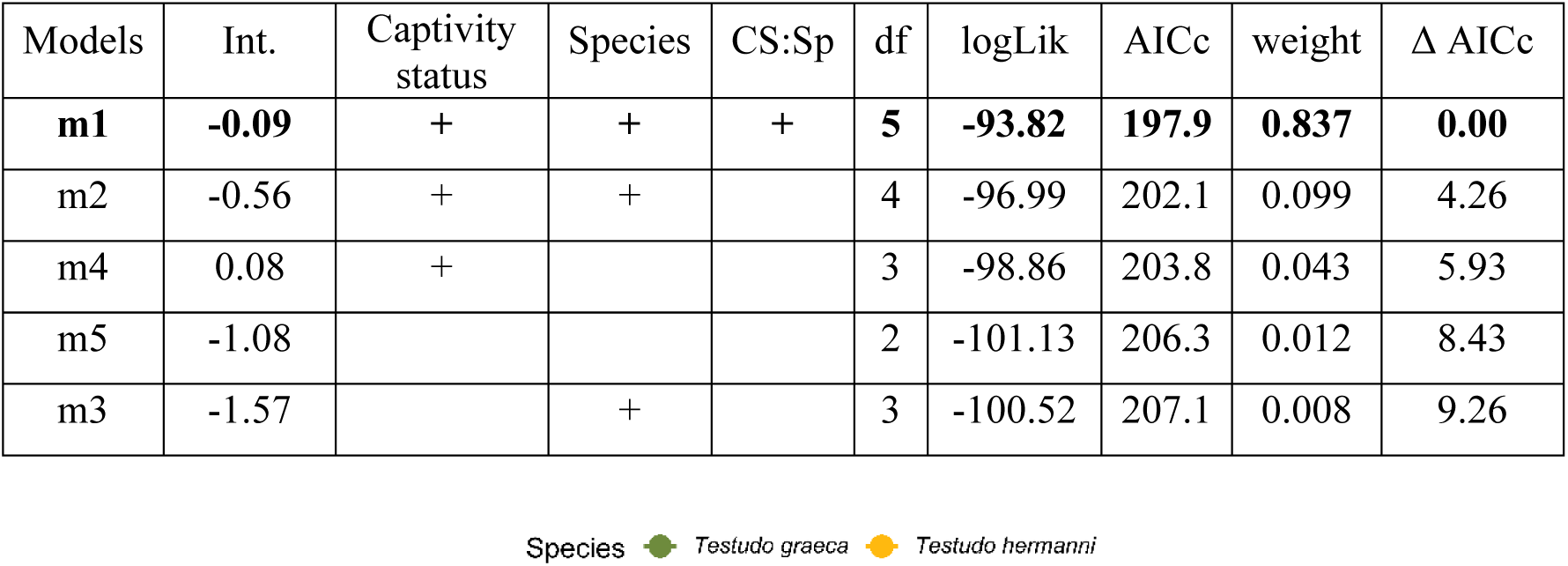
Model selection based on AICc strongly supported the model including captivity status, species, and their interaction (CS:Sp). All alternative models received considerably less support (ΔAICc > 4). Random terms in all models: (1 | sampling region).

| Models | Int. | Captivity status | Species | CS:Sp | df | logLik | AICc | weight | $\Delta AICc$ |
| --- | --- | --- | --- | --- | --- | --- | --- | --- | --- |
| **m1** | **-0.09** | + | + | + | **5** | **-93.82** | **197.9** | **0.837** | **0.00** |
| m2 | -0.56 | + | + |  | 4 | -96.99 | 202.1 | 0.099 | 4.26 |
| m4 | 0.08 | + |  |  | 3 | -98.86 | 203.8 | 0.043 | 5.93 |
| m5 | -1.08 |  |  |  | 2 | -101.13 | 206.3 | 0.012 | 8.43 |
| m3 | -1.57 |  | + |  | 3 | -100.52 | 207.1 | 0.008 | 9.26 |

## Discussion

This study presents the first assessment of *M. agassizii* prevalence in wild Mediterranean tortoise populations in Spain and contributes key epidemiological information on patterns of infection among *Testudo* species. Using samples obtained across multiple regions and including both wild and captive individuals, our results reveal marked differences in prevalence between captivity status and species. In particular, *T. graeca* showed very low prevalence in wild populations, contrasting with moderate to high prevalence in captive individuals, whereas *T. hermanni* showed comparatively higher prevalence in both wild and captive settings, with consistently elevated values in Catalonia. Across both species, infection was more frequently detected in captive tortoises, supporting the view that captivity may play an important role in pathogen maintenance and transmission. Together, these findings help to address the limited knowledge on *M. agassizii* occurrence in Mediterranean tortoises in Spain (Salinas et al. 2011b; Kolesnik et al. 2017; Martínez-Silvestre et al. 2021, 2025; Aranda et al. 2025), and provide a useful baseline for future evaluations of disease risk in threatened *Testudo* populations.

Our results reinforce the hypothesis that captive environments play a central role in shaping the epidemiology of *M. agassizii* in Europe, where high-density housing, mixed-species collections, and the pet trade may facilitate pathogen maintenance and dissemination (Origgi et al. 2004; Soares et al. 2004; Salinas et al. 2011b; Kolesnik et al. 2017; Galosi et al. 2023; Louro et al. 2025; Cacciotto et al. 2025). PCR-based surveys across Europe consistently report higher detection rates in captive tortoises than in free-ranging populations. In captive collections, reported prevalence generally range between 40% and 70% (Kolesnik 2017; Galosi et al. 2023; Aranda et al. 2025; Louro et al. 2025), with a single study documenting lower values around 6% (Salinas et al. 2011b). In contrast, wild populations typically show lower and more variable PCR detection rates, ranging from 0% to approximately 37% (Mathes 2003; Lecis et al. 2011; Ballouard et al. 2021). Similarly, in wild gopher tortoises in United States of America, PCR prevalence varies widely among localities, from absence of detection to very high local values (Jacobson et al. 2014; Weitzman et al. 2017), highlighting the spatial heterogeneity characteristic of natural systems. In this work, captive *T. graeca* from Alicante and Murcia exhibited particularly elevated infection levels, whereas prevalence was slightly lower in Mallorca. Similarly, captive *T. hermanni* displayed high infection rates across facilities. Notably, according to long-term observations from recovery centers, *T. hermanni* rarely experiences mortality during captivity, whereas *T. graeca* shows substantially higher mortality under similar conditions (Ferrández 2019). This contrast raises the possibility that *M. agassizii* infections may differentially affect clinical outcomes between species, with higher mortality or reduced resilience in *T. graeca* compared to *T. hermanni*. Targeted clinical and experimental studies are required to confirm species-specific effects. Indeed, high mortality registered in long-term *T. graeca* records in the Santa Faz (Alicante) and El Valle (Murcia) recovery centers (Saiz 2017) raises critical questions regarding the role of *M. agassizii* in disease severity and mortality for the species.

Coinfections with herpesviruses and other pathogens are commonly reported in tortoises infected with *M. agassizii* (Soares et al. 2004; Salinas et al. 2011b; Kolesnik et al. 2017; Ballouard et al. 2021). Combinations of high *Mycoplasmopsis* prevalence and viral coinfections may exacerbate morbidity and contribute to severe clinical manifestations (Jacobson et al. 2014; Aiello et al. 2018; Sandmeier et al. 2022). However, testing for additional pathogens was not carried out in the present study, but would be of interest in future evaluations of the impact of infections on mortality in these species.

The detection of *M. agassizii* in a wide range of Testudinidae across Europe, North America, North Africa, and even Asia underscores the pathogen’s global distribution (Brown et al. 1999; Berry et al. 2015; Kolesnik et al. 2017; Galosi et al. 2023). In this context, given the consistently higher infection probability in captivity, the movement of captive tortoises, whether through legal or illegal animal trade may constitute a key driver of *M. agassizii* infection dynamics (Lecis et al. 2013; Ballouard et al. 2021; Rush et al. 2021; Apakupakul et al. 2025). Support for this interpretation comes from studies showing higher prevalence in wild tortoise populations located near urbanised areas compared to more remote sites (Berry et al., 2014), suggesting pathogen spillover associated with anthropogenic activities. Moreover, recent genetic analyses have revealed high strain diversity and limited geographic structuring among captive tortoises in southern Europe (Cacciotto et al., 2025), further indicating that animal movement and mixing in managed environments may facilitate both dissemination and genetic diversification of *M. agassizii*. More broadly, recent evidence indicates that emerging infectious diseases in chelonians may exhibit synergistic interactions with global trade and habitat alteration, producing stronger combined effects than would be expected from each threat acting independently (Rodríguez-Caro et al., 2025). In this framework, pathogen dynamics in Mediterranean tortoises cannot be interpreted on an individual basis, but rather as part of a network of interacting anthropogenic pressures.

The absence of *M. agassizii* in wild *T. graeca* in mainland Spain is striking, particularly given the species’ broad distribution and the sampling effort in Almería. Only a single wild individual from Mallorca tested positive; this individual also exhibited poor body condition and oral mucosal pallor at the time of sampling. This case was detected at a locality assumed to host only *T. graeca*, where the presence of *T. hermanni* was also recorded, consistent with the broader regional context of frequent human-mediated faunal introductions (Pinya et al. 2007; Silva-Rocha et al. 2018). Several non-exclusive processes may explain this pattern. One possibility is that wild *T. graeca* populations in the south-eastern Iberian Peninsula have experienced little or no historical exposure to *M. agassizii,* potentially due to their geographic isolation and limited opportunities for pathogen introduction (Graciá et al. 2013, 2020). Alternatively, infections may occur only sporadically and fail to persist because chronically infected hosts contribute unevenly to transmission, owing to variable pathogen shedding and immune responses. In addition, ecological factors such as low population densities, reduced contact rates, or habitat structure may further constrain pathogen transmission (Aiello et al. 2014, 2018; Jacobson et al. 2014; Rodríguez-Caro et al. 2016). Together, these findings suggest that *T. graeca* populations in Spain may be immunologically naive, rendering them particularly vulnerable to pathogen introduction from external sources.

In contrast, wild *T. hermanni* exhibited high prevalence in north-eastern Iberian Peninsula, with values comparable to those reported in southern France (Ballouard et al. 2021), suggesting that *M. agassizii* is well established in natural populations of *T. hermanni* within this portion of its range. Long-term monitoring indicates that some populations, such as those in Albera, have persisted for decades while maintaining moderate to relatively high densities despite documented population declines and demographic imbalances (Bertolero et al. 2020; Mascort 2025). This apparent capacity to persist in the presence of infection may reflect species-specific differences in susceptibility or immune response, influencing transmission dynamics. The lower prevalence detected in Mallorca may be influenced by limited sample size, which could have reduced the probability of detecting infected individuals. Overall, the contrasting epidemiological *M. agassizii* profiles in *T. graeca* and *T. hermanni* highlight fundamental interspecific differences in host–pathogen interactions that merit further investigation.

The contrasting prevalence patterns between species, together with the concentration of infections in captive reservoirs, have direct implications for conservation management (Aiello et al., 2014; Warne and Chaber 2023). Our results suggest that wild *T. graeca* populations, in particular, may lack prior exposure to *M. agassizii*, making them potentially vulnerable to high-impact outbreaks if the pathogen is introduced from captive sources. Preventing the release of captive individuals into the wild, implementing rigorous health screening and pathogen surveillance for individuals bred or maintained in captivity prior to any conservation translocation, and strengthening biosecurity measures in wildlife recovery centers are essential actions to minimize pathogen spillover and support long-term conservation efforts.

## Author contribution

MCB, AG and EG contributed to the study conception and design. MCB, AG, AMS, JB, BSF, RRC and EG participated in data collection. Laboratory analysis were performed by RM. Data processing and analysis were performed by MCB. EG and AG were responsible for securing and managing the funding that supported this study and for supervising MC, as this work was conducted as part of her doctoral thesis. First draft of the manuscript was written by MCB and all authors commented on previous versions of the manuscript. All authors read and approved the final manuscript.

## Statement on welfare of animals

All procedures involving animals followed all institutional and national guidelines for the care and use of animals and were approved by the Miguel Hernández University Research Ethics Committee (UMH.DBA.AGC.01.22).

## Data availability statement

All data and code supporting this study are available at the Zenodo Digital Repository: 10.5281/zenodo.18803824

## Acknowledgments

We thank the Doñana Biological Reserve (DBR-ICTS), Doñana National Park, and the Junta de Andalucía for granting the necessary permits to conduct this study. We are grateful to the Consorci per a la Recuperació de la Fauna de les Illes Balears (COFIB) for their valuable assistance during sampling activities in Mallorca, and to the Govern de les Illes Balears for the corresponding authorizations. We also acknowledge the CRARC (Centre de Recuperació d’Amfibis i Rèptils de Catalunya), the CRT (Centre de Recuperació de Tortugues de Catalunya), the Valle Recovery Center, and the Santa Faz Wildlife Recovery Center for facilitating access to animals and biological samples. We thank the staff of Biocyma for their collaboration during field sampling in Almería. Additional permits were granted by the Junta de Andalucía and the Región de Murcia.

## References

Adamovicz L, Allender MC, Gibbons PM (2020) Emerging infectious diseases of chelonians: An update. Vet Clin North Am Exot Anim Pract 23(2):263–283. 10.1016/j.cvex.2020.01.014

Aiello CM, Esque TC, Nussear KE, Emblidge PG, Hudson PJ (2019) The slow dynamics of mycoplasma infections in a tortoise host reveal heterogeneity pertinent to pathogen transmission and monitoring. Epidemiol Infect 147: e12. 10.1017/S0950268818002613

Aiello CM, Nussear KE, Walde AD, Esque TC, Emblidge PG, Sah P, Bansal S, Hudson PJ (2014) Disease dynamics during wildlife translocations: disruptions to the host population and potential consequences for transmission in desert tortoise contact networks. Anim Conserv 17:27–39. 10.1111/acv.12147

Altizer S, Harvell D, Friedle E (2003) Rapid evolutionary dynamics and disease threats to biodiversity. Trends Ecol Evol 18(11):589–596. 10.1016/J.TREE.2003.08.013

Aranda C, Heredia A, Esperón F (2025) Clinical Trial of Mycoplasma treatment in captive Testudo graeca populations. In: Graciá, E., Rodríguez-Caro, R.C., & Giménez, A. (eds.). Conservation of Mediterranean tortoises under global change. Madrid. Asociación Herpetológica Española.

Ballouard JM, Bonnet X, Jourdan J, Martínez-Silvestre A, Gagno S, Fertard B, Caron S (2021) First detection of herpesvirus and prevalence of mycoplasma infection in free-ranging Hermann’s tortoises (*Testudo hermanni*), and in potential pet vectors. Peer Community J 2. 10.1101/2021.01.22.427726

Berry KH, Brown MB, Vaughn M, Gowan TA, Hasskamp MA, Torres MCM (2015) Mycoplasma agassizii in Morafka’s desert tortoise (*Gopherus morafkai*) in Mexico. J Wildl Dis, 51(1):89–100. 10.7589/2014-04-083

Bertolero A, Budó J, Torres N (2020) Característiques poblacionals de la tortuga mediterrània Testudo hermanni hermanni Gmelin a la serra de l’Albera (Reptilia: Testudinidae). Butl Inst Catalana Hist Nat 84:131–136.

Bertolero A, Pretus JL (2015) La tortuga mediterránea (*Testudo hermanni*) en las islas Baleares. BAHE 26(2): 43–46.

Brown D, Crenshaw B, McLaughlin G, Schumacher I, McKenna C, Klein P, et al. (1995) Taxonomic analysis of the tortoise mycoplasmas *Mycoplasma agassizii* and Mycoplasma testudinis by 16S rRNA gene sequence comparison. Int J Syst Bacteriol 45(2):348–50. 10.1099/00207713-45-2-348

Brown MB, McLaughlin GS, Klein PA, Crenshaw BC, Schumacher IM, Brown DR Jacobson ER (1999) Upper respiratory tract disease in the Gopher tortoise is caused by *Mycoplasma agassizii*. J Clin Microbiol 37: 2262–2269. 10.1128/jcm.37.7.2262-2269.1999

Cacciotto C, Zobba R, Louro M, Alves M, Valença A, Patrício R, Bazzoni E, Pittau M, Lecis R, Alberti A (2025) Mycoplasma agassizii Multilocus Sequence Typing Using Nanopore Sequencing: Insights Into Genetic Diversity and Isolate Characterization. Ann N Y Acad Sci 1554(1): 153–162. 10.1111/nyas.70128

Canós-Burguete M, Giménez A, Martínez-Silvestre A, Budó J, Marschang RE, Sánchez-Ferreiro B, Rodríguez-Caro RC, Graciá E (2026) Mycoplasmopsis agassizii prevalence in mediterranean tortoises (Testudo hermanni and Testudo graeca) [Data set]. Zenodo. 10.5281/zenodo.18803824

Chen C, Wang J, Holyoak M, Lin L, Wang Y (2025) Global assessment of current extinction risks and future challenges for turtles and tortoises. Nat Commun 16(1):7114. 10.1038/s41467-025-62441-2

Daszak P, Cunningham AA, Hyatt AD (2000) Emerging Infectious Diseases of Wildlife Threats to Biodiversity and Human Health. Science 287:443–449. 10.1126/science.287.5452.443

Di Nicola MR, Rubiola S, Cerullo A, Basciu A, Massone C, Zabbia T, Dorne JLCM, Acutis PL, Marini D (2025) Microorganisms in wild European reptiles: bridging gaps in neglected conditions to inform disease ecology research. Int J Parasitol Parasites Wildl 101113. 10.1016/j.ijppaw.2025.101113

Drake KK, Aiello CM, Bowen L, Lewison RL, Esque TC, Nussear KE, Waters SC, Hudson PJ (2019) Complex immune responses and molecular reactions to pathogens and disease in a desert reptile (*Gopherus agassizii*). Ecol Evol 9(5):2516–2534. 10.1002/ece3.4897

Faulhaber MM, Tardy F, Saul F, Müller E, Pees M, Marschang RE. (2025) Detection of *Mycoplasma* spp. from snakes from five different families. BMC Vet Res 21(1):38. 10.1186/s12917-025-04487-4

Ferrández M (2019) Management of captive populations of *Testudo graeca* in the Fauna Recovery Center of Santa Faz (Alicante). In: Mediterranean Workshop to Develop Tortoise Conservation Strategies, Alicante, Spain, 27–29 October 2019.

Field EK, Hartzheim A, Terry J, Dawson G, Haydt N, Neuman-Lee LA (2022) Reptilian innate immunology and ecoimmunology: what do we know and where are we going? Integr Comp Biol 62(6):1557–1571. 10.1093/icb/icac116

Galosi L, Ridolfi N, Fellini C, Pelizzone I, Cusaro S, Marchetti G, Canonico M, Ghefi E, Di Girolamo N, Preziuso S (2023) Detection and identification of *Mycoplasmopsis agassizii* in captive tortoises with different clinical signs in Italy. Animals, 13(4):588. 10.3390/ani13040588

Graciá E, Giménez A, Anadón JD, Botella F, García-Martínez S, Marín M (2011) Genetic patterns of a range expansion: the spur-thighed tortoise *Testudo graeca graeca* in Southeastern Spain. AMRE, 32(1):49–61. 10.1163/017353710X542985

Graciá E, Giménez A, Anadón JD., Harris DJ, Fritz U, Botella F (2013) The uncertainty of Late Pleistocene range expansions in the western Mediterranean: a case study of the colonization of south-eastern Spain by the spur-thighed tortoise, *Testudo graeca*. J Biogeogr 40: 323–334. 10.1111/jbi.12012

Graciá E, Rodríguez-Caro RC, Andreu AC, Fritz U, Giménez A, Botella F (2017a). Human-mediated secondary contact of two tortoise lineages results in sex-biased introgression. Sci Rep 7(1):4019. 10.1038/s41598-017-04208-4

Graciá, E, Rodríguez-Caro RC, Ferrández M, Martínez-Silvestre A, Pérez-Ibarra I, Amahjour R, Aranda C, Aissa Benelka-Di H, Bertolero A, Biaggini M, Botella F, Budó J, Cade-Nas V, Chergui B, Corti C, Esperón F, Esteve-Selma MÁ, Fahd S, García de La Fuente MI, Golubovic A, Heredia A, Jiménez-Franco MV, Arakelyan M, Marini D, Martínez-Fernández J, Martínez-Pastor MC, Mascort R, Mira-Jover A, Pascual-Rico R, Perera-Leg A, Pfau B, Pinya S, Santos X, Segura A, Semaha MMJ, Soler-Massana J, Vidal JM, Giménez A (2020). From troubles to solutions: Conservation of mediterranean tortoises under global change. Basic Appl Herpetol 34:5–16. 10.11160/bah.196

Graciá E, Vargas-Ramírez M, Delfino M, Anadón JD, Giménez A, Fahd S, Corti C, Jdeidi TB, Fritz U (2017b) Expansion after expansion: dissecting the phylogeography of the widely distributed spur-thighed tortoise, *Testudo graeca* (Testudines: Testudinidae). Biol J Linn Soc 121(3):641–654. 10.1093/biolinnean/blx007

Hidalgo-Vila J, Martínez-Silvestre A, Pérez-Santigosa N, León-Vizcaíno L, Díaz-Paniagua C (2020) High prevalence of diseases in two invasive populations of red-eared sliders (*Trachemys scripta elegans*) in southwestern Spain. AMRE, 41(4), 509–518. 10.1163/15685381-bja10021

Iglesias R, García-Estévez JM, Ayres C, Acuña A, Cordero-Rivera A (2015) First reported outbreak of severe spirorchiidiasis in Emys orbicularis, probably resulting from a parasite spillover event. Dis Aquat Org 113:75–80. 10.3354/dao02812

Jacobson ER, Brown MB, Wendland LD, Brown DR, Klein PA, Christopher MM, Berry KH (2014) *Mycoplasmosis* and upper respiratory tract disease of tortoises: a review and update. Vet J 201(3):257–64. 10.1016/j.tvjl.2014.05.039

Kolesnik E, Obiegala A, Marschang R (2017) Detection of *Mycoplasma* spp., herpesviruses, topiviruses, and ferlaviruses in samples from chelonians in Europe. J Vet Diagn Invest 29(6): 820–832. 10.1177/104063871772238

Lecis R, Paglietti B, Rubino S, Are BM, Muzzeddu M, Berlinguer F, Chesa B, Pittau M, Alberti A (2011) Detection and characterization of Mycoplasma spp. and Salmonella spp. in free-living European tortoises (*Testudo hermanni*, Testudo graeca, and Testudo marginata). J Wildl Dis 47(3):717–724. 10.7589/0090-3558-47.3.717

Louro M, Patrício R, Pereira A, Valença A, Alves M (2025) First molecular detection of *Mycoplasma agassizii* in captive tortoises in Portugal. Front Vet Sci 12:1652362. 10.3389/fvets.2025.1652362

Luiselli L, Starita A, Carpaneto GM, Segniagbeto GH, Amori G (2016) A short review of the international trade of wild tortoises and freshwater turtles across the world and throughout two decades. Chelonian Conserv Biol 15(2):167–172. 10.2744/CCB-1216.1

Marenzoni ML, Stefanetti V, Del Rossi E, Zicavo A, Scuota S, Origgi FC, Deli G, Corti C, Marinucci MT, Oliveri O (2022) Detection of Testudinid alphaherpesvirus, *Chlamydia* spp., *Mycoplasma* spp., and *Salmonella* spp. in free-ranging and rescued Italian *Testudo hermanni hermanni*. Vet Ital, 58(1), 25–34. 10.12834/VetIt.1915.13833.1

Martínez-Silvestre A, Soler J, Iturria D, Geisler G, Marchang RE (2022) Estado sanitario de una población introducida de tortuga mediterránea (*Testudo hermanni*) en la Sierra del Montsant (Tarragona). BAHE 33:129–131.

Martínez-Silvestre A (2025) Emerging diseases and biosecurity in the conservation of Mediterranean Tortoises *Testudo graeca* and *T. hermanni**. In*: Graciá, E., Rodríguez-Caro, R.C., & Giménez, A. (eds.). Conservation of Mediterranean tortoises under global change. Madrid. Asociación Herpetológica Española.

Mascort R (2025) Long-term monitoring of a population core of western Hermann’s tortoise, *Testudo hermanni hermanni*, at Serra de l’Albera, in the north-eastern Iberian Peninsula. *In*: Graciá, E., Rodríguez-Caro, R.C., & Giménez, A. (eds.). Conservation of Mediterranean tortoises under global change. Madrid. Asociación Herpetológica Española.

Mateo JA, Oliver JA, Mayol J (2011) Las translocaciones de tortugas de tierra en Mallorca, treinta años de manejo y liberaciones. La Conservación de las Tortugas de Tierra en España, pp. 51–56. Conselleria de Medi Ambient i Mobilitat, Govern de les Illes Balears, Palma.

Mathes KA (2003). Untersuchungen zum Vorkommen von Mykoplasmen und Herpesviren bei freilebenden und in Gefangenschaft gehaltenen Mediterranen Landschildkröten (Testudo hermanni, Testudo graeca graeca und Testudo graeca ibera) in Frankreich und Marokko. VVB Laufersweiler.

Origgi FC, Romero CH, Bloom DC, Klein, PA, Gaskin JM, Tucker SJ, Jacobson ER (2004) Experimental transmission of a herpesvirus in Greek tortoises (*Testudo graeca*). Vet Pathol 41(1):50–61. 10.1354/vp.41-1-50

Pfau B, Budó J, Martínez-Silvestre A (2020) Autochthone und allochthone Landschildkröten in Katalonien, Update. Testudo (SIGS) 29(3).

Pinya S (2011) Situación actual de la Tortuga Mora (*Testudo graeca* L.) en le Isla de Mallorca. En J.A. Mateo (ed.). La Conservación de las Tortugas de Tierra en España, pp. 7–12. Conselleria de Medi Ambient i Mobilitat, Govern de les Illes Balears, Palma.

Rios FM, Zimmerman LM (2015) Immunology of reptiles. eLS, 1–7.

Rodríguez-Caro RC, Lima M, Anadón JD, Graciá E, Giménez A (2016). Density dependence, climate and fires determine population fluctuations of the spur-thighed tortoise *Testudo graeca*. J Zool 300(4):265–273. 10.1111/jzo.12379

Rodríguez-Caro RC, Graciá E, Blomberg SP, Cayuela H, Carmona CP, Pérez-Mendoza HA, Giménez, A, Salguero-Gómez R (2023) Anthropogenic impacts on threatened species erode functional diversity in chelonians and crocodilians. Nat Commu 14, 1542. 10.1038/s41467-023-37089-5

Rodríguez-Caro RC, Gumbs R, Graciá E, Blomberg SP, Cayuela H, Grace MK, Salguero-Gómez R (2025) Synergistic and Additive Effects of Multiple Threats Erode Phylogenetic and Life History Strategy Diversity in Testudines and Crocodilia. Ecol Lett, 28(6), e70147. 10.1111/ele.70147

Rush ER, Dale E, Aguirre AA (2021). Illegal wildlife trade and emerging infectious diseases: Pervasive impacts to species, ecosystems and human health. Animals, 11(6), 1821. 10.3390/ani11061821

Salinas M, Altet L, Clavel C et al. (2011a) Genetic assessment, illegal trafficking and management of the Mediterranean spur-thighed tortoise in Southern Spain and Northern Africa. Conserv Genet 12:1–13. 10.1007/s10592-009-9982-1

Salinas M., Francino O, Sánchez A, Altet L (2011b). *Mycoplasma* and Herpesvirus PCR Detection in Tortoises with Rhinitis-stomatitis Complex in Spain. J Wildl Dis 47: 195–200. 10.7589/0090-3558-47.1.195

Sandmeier FC, Tracy CR, Hunter K (2009) Upper respiratory tract disease (URTD) as a threat to desert tortoise populations: A reevaluation. Biol Conserv 142(7):1255–1268. 10.1016/j.biocon.2009.02.001

Sandmeier FC, Leonard KL, Weitzman CL, Tracy CR (2022) Potential facilitation between a commensal and a pathogenic microbe in a wildlife disease. EH 19(3):427–438. 10.1007/s10393-022-01603-w

Schönbächler K, Segner H, Amphimaque B, Friker B, Hofer A, Lange B, Stirn M, Pantchev N, Origgi FC, Hoby S (2022) Health assessment of captive and free-living european pond turtles (*Emys orbicularis*) in Switzerland. J Zoo Wildl Med 53(1):159–172. 10.1638/2020-0117

Segura A, Delibes-Mateos M, Acevedo P (2020) Implications for conservation of collection of Mediterranean spur-thighed tortoise as pets in Morocco: Residents’ perceptions, habits, and knowledge. Animals, 10(2), 265. 10.3390/ani10020265

Semaha MJ, Rodríguez-Caro RC, Giménez A, Fahd S, Graciá E (2025) Captive-introduced tortoises in wild populations: can we identify them by shell morphology?. Eur J Wildl Res, 71(1):13. 10.1007/s10344-024-01893-1

Shearer I, Türkozan O (2024). Global Testudo Trade: Update and Recent Trends. Chelonian Conserv Biol 23(2):161–168. 10.2744/CCB-1643.1

Silva-Rocha I, Montes E, Salvi D, Sillero N, Mateo JA, Ayllón E, Pleguezuelos JM, Carretero MA (2018) Herpetological history of the Balearic Islands: when aliens conquered these islands and what to do next. In Histories of Bioinvasions in the Mediterranean (pp. 105–131). Cham: Springer International Publishing.

Soares JF, Chalker VJ, Erles K, Holtby S, Waters M, McArthur S (2004) Prevalence of Mycoplasma agassizii and chelonian herpesvirus in captive tortoises (*Testudo* sp.) in the United Kingdom. J Zoo Wildl Med 35(1):25–33. 10.1638/02-092

Warne RK, Chaber AL (2023) Assessing disease risks in wildlife translocation projects: a comprehensive review of disease incidents. Animals 13(21):3379. 10.3390/ani13213379

Weitzman CL, Sandmeier FC, Tracy CR (2017). Prevalence and diversity of the upper respiratory pathogen Mycoplasma agassizii in Mojave desert tortoises (*Gopherus agassizii*). Herpetologica 73(2):113–120. 10.1655/Herpetologica-D-16-00079.1

